# Group Similarity Constraint Functional Brain Network Estimation for Mild Congititive Impairment Classification

**DOI:** 10.1101/734574

**Authors:** Xin Gao, Xiaowen Xu, Weikai Li, Rui Li

**Affiliations:** Shanghai Universal Medical Imaging Diagnostic Center, Shanghai 20030, China; Tongji University School of Medicine, Tongji University, Shanghai 200092, China; Department of Medical Imaging, Tongji Hospital, Shanghai 200092, China; College of Information Science and Engineering, Chongqing Jiaotong University, Chongqing 400074, China; School of mechanical and precision instrument engineering, Xi ‘an university of technology, Xi’an 710048, China

**Keywords:** Functional Brain Network, Functional Magnetic Resonance Imaging, Group Constraint, Mild cognitive impairment (MCI), Pearson’s correlation, Partial correlation, Low-rank Regularizer

## Abstract

Functional brain network (FBN) provides an effective biomarker for understanding brain activation patterns, which also improve the diagnostic criteria for neurodegenerative diseases or the information transmission of brain. Unfortunately, despite its efficiency, FBN still suffers several challenges for accurately estimate the biological meaningful or discriminative FBNs, under the poor quality of functional magnetic resonance imaging (fMRI) data as well as the limited understanding of human brain. Hence, there still a motivation to alleviate those issues above, it is currently still an open field to discover. In this paper, a novel FBN estimation model based on group similarity constraints is proposed. In particular, we extend the FBN estimation model to the tensor form and incorporate the trace-norm regularizer for formulating the group similarity constraint. In order to verify the proposed method, we conduct experiments on identifying Mild Cognitive Impairments (MCIs) from normal controls (NCs) based on the estimated FBNs. The experimental results illustrated that the proposed method can construct a more discriminative brain network. Consequently, we achieved an 91.97% classification accuracy which outperforms the baseline methods. The *post hoc* analysis further shown more biologically meaningful functional brain connections obtained by our proposed method.

## 1. Introduction

As neurodegenerative disorders, Alzheimer’s disease (AD) is one of the most common cause of dementia (Gaugler et al. 2016). According to a recent report (Bain et al. 2008), the incidence of AD doubles per 5 year after the age 6. The AD seriously interferes the human daily life, affects memory and ability to reason and communication, and eventually causes deaths. Unfortunately, so far, there still no effective treatment for AD. Therefore, it is quite important for delaying the onset and progression of AD during its early stage based on the pharmacological or behavioral interventions.

Mild cognitive impairment (MCI) is often considered to be a critical time window and treatment period for the prediction or delaying the conversion in AD (Wee et al. 2012a). In some recent statistical studies, nearly 10-15% patients with MCI each year tend to develop probable AD (Grundman et al. 2004, Misra, Fan and Davatzikos 2009). The early detection and accurate diagnosis for MCI is considered to be a significant means of slowing down the progression of AD (Association 2017).

As a successful non-invasive technique, functional magnetic resonance imaging (fMRI) provides an effective measurement for revealing the brain activity or pattern (Brunetti et al. 2006, Jin et al. 2010, Kevin et al. 2008). However, due to its spontaneous brain activities are random and asynchronous across subjects or scanners, it is still challengeable for identifying patients from normal controls (NC). Furthermore, the high-order statistical information based FBN served as a new perspective for discovering the brain activity and connection pattern, which improved the stability for understanding brain information (Rosa et al. 2015, Smith et al. 2011b, Sporns 2011, Stam 2014, Wee et al. 2012b). Besides, series of researches illustrate that the functional brain network are highly related to some neurological or psychological diseases, such as AD (Huang et al. 2009, Liu et al. 2012, Supekar et al. 2008), MCI (Fan and Browndyke 2010, Wee et al. 2012b, Wee et al. 2014, Yu et al. 2016), autism spectrum disorder (ASD) (Gotts et al. 2012, Theije et al. 2011), parkinsonism disease (PD) (Baggio et al. 2014) and so on. Note that, all of these depend heavily on the quality of the final estimated FBN, so, it is crucial for estimating the more reliable FBNs (Li et al. 2019a).

The second-order statistics (i.e., correlation) based FBN estimation methods play a dominant role in FBN estimation. According to a FBN research review (Smith et al. 2011a), the correlation based methods such as Pearson’s correlation (PC) (Li et al. 2017), sparse representation (SR) (Lee et al. 2011, Zhou, Wang and Ogunbona 2014), are generally worked more sensitive than complex high-order methods. However, due to the influence of noise exists in the observed data, the correlation based brain networks will inevitably have dense connections and thus contain a lot of noise or false connections. One solution is to introduce sparse priors, such as the thresholding method or the SR (LASSO) method. Actually, the topological structure of FBN is no more than just sparsity (Sporns 2011). Therefore, several studies are focused on incorporating more biological priors of FBN towards more discriminative FBNs. In practice, the commonly-used priors include sparsity, modularity, group-sparsity, low-rank and scale-free (Lee et al. 2011, Li et al. 2017, Qiao et al. 2016b, Wee, Yap and Shen 2016, Yu et al. 2016). Moreover, the priors can also obtained from the data quality (Li et al. 2019a) and other high quality data (Li et al. 2019b). Note that, most of them can be summarized into the regularized framework, which illustrate that a reliable FBN estimation model should not only fit the data well, but also effectively encode priors of the brain organization (Qiao et al. 2016b).

However, most existing FBN estimation methods mainly focused on a single participant, which rarely consider the inter-group information cross participants. Due to the limited data quality, the FBNs estimated by these methods are easily tend to have poor performance. This is attributed to the human brain networks have a group similarity prior (Wee et al. 2014), while these methods ignoring the relations inter group. In addition, it will result in different network topological structures across subjects, and thus inevitably make the comparison between subjects difficult and thus possibly degrade generalization performance of trained classifiers. Besides, recent studies have also illustrated that group constraints can effectively improve the performance of estimated FBNs (Wee et al. 2014, Yu et al. 2016). The existing method for group constraints mainly based on the penalty of group sparsity (i.e., *l*_2,1_-norm) for mitigating the inter-subject variability. However, it is strong penalty to constraint all of the estimated FBN, which will cause the samples of different groups (e.g., MCI and NC) to interact with each other, due to the additional *l*_2_-norm penalization. Therefore, this approach may hinder performance improvements.

In this paper, based on the regularization framework, we try to incorporate the group similarity constraint into the FBN estimation model and relax the *l*_2,1_-norm penalty. In particular, we first extended the traditional matrix regularization framework to the tensor regularization framework for obtaining group similar information. Then, we formulate the group similarity prior as a tensor low rank regularizer and incorporate it into the FBN estimation model. Since the low-rank is NP-hard, we optimize its upper limit (i.e. trace norm regularizer) for better calculating efficiency. In particular, we adopt PARAFAC decomposition to calculate its eigenvalues (Liu, Bourennane and Fossati 2012), and design a proximal operator to estimate the FBN with group similarity constraint. In the end, we incorporate the trace norm regularizer into the SR and PC model as a simple test platform. For verifying the proposed methods, we adopted estimated FBN for MCI identification. In fact, the proposed method uses the group similarity constraints to shrink the solution space of the FBN, thus can estimate the more discriminative FBNs effectively. The highlight of this paper is given as follows:

1. Compared with the traditional FBN estimation methods of estimating FBN for a single participant, we proposed a group FBN estimation scheme, which builds the whole FBN simultaneously. In this way, The FBNs with intra-group priors can be naturally incorporated by the tensor regularization.
2. We incorporate the group similarity constraint into the FBN estimation model by a low rank regularizer. in addition, we further relaxed it into a trace norm regularizer and design an optimization algorithm for estimating the group similarity based FBNs.
3. We use the group similarity based FBN for identifying the MCIs from NCs. The experimental results show that the proposed scheme can achieve the 81.52% classification accuracy, which outperforms the baseline methods. Moreover, the provides methods can provide more biological meaningful connections.
4. We provide a FBN estimation module for modeling the group similarity prior, which is flexible for incorporating into other FBN estimation model. The experimental results show that the proposed module can effectively improve the accuracy of the estimated FBNs for MCI classification.
5. We identified the most significant functional connections and the most discriminative brain regions based on the proposed FBN estimation model. The analysis of functional connectivity and graph theory attributes can be used to discovering the biological meaningful biomarkers and further elucidate the topological properties of brain network in MCI.

The remainder of this paper is organized as follows. In Section 2, we introduce the material and methods which are selected in this paper. Specifically, we first introduce the data acquisition and briefly review the two most related methods, i.e., PC and SR. Then, we give the proposed methods i.e. group similarity based FBN estimation scheme (TLR), including the motivations, models and algorithms for these two methods. In Section 3, we evaluate the proposed methods with experiments on identifying MCI. In Section 4, we discuss our findings and prospects of our work. In Section 5, we conclude the entire paper briefly.

## 2. Material and Methods

### 2.1 Data Acquisition

In this study, we adopt the publically available neuroimaging data from the Alzheimer’s Disease Neuroimaging Initiative (ADNI) database (Jack et al. 2010)^1^. ANDI was launched in 2003 by the National Institute on Aging, the National Institute of Biomedical Imaging and Bioengineering, the Food and Drug Administration, private pharmaceutical companies and nonprofit organizations. Initially, the goal of ADNI is to define biomarkers for use in clinical trials and to determine the best way to measure the treatment effects of AD therapeutics.

In particular, 137 participants, including 68 MCIs and 69 NCs, are adopted in this experiment, which is also similar as (Zhou et al. 2018). The scanning parameter includes: TR/TE = 3000/30mm, flip angle = 80, imaging matrix=64×64, 48 slices, 140 volumes, and voxel thickness = 3.3mm. SPM8 toolbox2 and DPARSFA (version 2.2) (Chao-Gan and Yu-Feng 2010) are used to preprocess the fMRI data according to the well accepted pipeline. The preprocessing pipeline includes remover first 10 volumes, Slice timing, Realign, Normalize, Spatially smooth, Temporally Detrend, Regression out covariates and Temporally filerting. For alleviating the head motion effect and artifacts, we follow the previous work (Chen et al. 2016, Chen et al. 2017), and exclude the subjects with more than 2.5 min (50 frames) data of FD>0.5 from further analysis (Power et al. 2012). Finally, depending on the automated anatomical labeling (AAL) atlas (Tzourio-Mazoyer et al. 2002), the pre-processed BOLD time series signals are partitioned into 116 ROIs. At last, we put these time series into a data matrix X ∈ R^137×116^. For more details, please refer to (Zhou et al. 2018).

### 2.2 Functional Brain Network Estimation

After obtaining the fMRI data matrix *X* from the R-fMRI data, the subsequent task is the FBN estimation. As we mentioned above, the correlation based FBN estimation methods have been demonstrated to be more sensitive than some complex higher-order methods (Smith et al. 2011b). Therefore, in this paper, we focus on the correlation-based methods and adopt it as a baseline. For better notation, we first define the data matrix (i.e., BOLD signal matrix) **X** ∈ *R^T×N^*, where T is the number of volumes and N is the number of ROIs. The fMRI time series associated with the ith ROI is represented by **x**_i_ ∈ *R^T^*, *i* = 1,…, *N.* A

#### 2. 2. 1 Correlation based Methods

As the most simplest and widely used FBN estimation scheme, Pearson’s Correlation (PC) based FBN estimation methods account for a large proportion in the study of FBNs (Smith et al. 2013). The edge weights of the FBN **W** = (*W_ij_*) ∈ *R^N×N^* can be calculated by PC as follows:

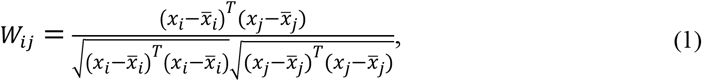

In Eq. (1), 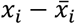 is a centralized counterpart of *x_i_*. Due to the effect of the noises mixed in the fMRI data, PC always generates dense FBNs. Thus, a thresholding scheme is often selected for sparsifying the PC-based FBNs, which aims for filtering out the noisy or weak connections. The PC based FBN can be expressed as follows:

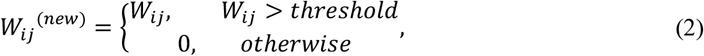

where *W_ij_*^(*new*)^ denotes the connection value between nodes i and j after thresholding.

Compared with PC measures the full correlation cross ROIs, the interaction among multiple ROIs is neglect due to the cofounding effect. In contrast, the partial correlation is proposed by regressing out the confounding effects from other ROIs. However, the partial correlation-based methods can be easily ill-posed due to the involvement of inverting the covariance matrix **Σ = X^T^X**. A base solution is to incorporate an *l*_1_-norm regularizer into the partial correlation model (Lee et al. 2011), which also naturally incroporates the sparsity prior (SR) of FBN. The model of SR is shown as follows:

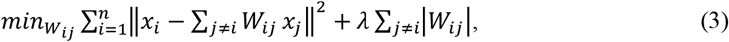

the matrix form is proposed as follows:

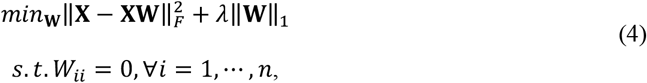

Note that the *l*_1_-norm regularizer in Eq. (4) plays a key role in achieving a sparse and stable solution (Lee et al. 2011).

#### 2. 2. 2 Regularization Framework for FBN estimation

According to the above description, both PC- and SR-based FBN estimation models can be summarized into the regularized FBN learning framework. We can naturally incorporate a regularized term and statistical information into the objective function for constructing a new platform to estimate FBNs. More specifically, the platform can be formulated using a matrix-regularized learning framework as follows:

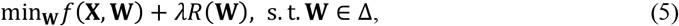

where *f*(**X, W**) models the statistical information of FBN, and *R*(**W**) is the regularization term for incorporating biological priors of FBN and stabilizing the solutions. In addition, some specific constraints such as symmetry or positive semi-definiteness may be included in Δ for shrinking the search space of **W**, which provides an effective way for obtaining a better FBN. The *λ* is a hyper-parameter for controling the balance between the first (data-fitting) term and the second (regularization) term.

In fact, most of the recently-proposed FBN estimation models (Higgins, Kundu and Guo 2018, Li, Zhu and Fan 2018, Wang et al. 2018, Zhou et al. 2018) can be unified under this regularized framework with different design of the two terms in Eq. (5). The popular data-fitting terms include 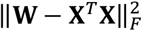 used in Eq. (2) and 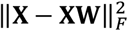 used in Eq. (4), while the popular regularization terms include l1-norm (Huang et al. 2010), trace norm and their combination (Qiao et al. 2016a), etc.

#### 2. 2. 4 Sparse and Low-rank based FBN estimation

Before we introduce the proposed method, we would like to brief review the sparse and low-rank based FBN estimation model (Qiao et al. 2016b). The sparsity and low-rank regularizer (SLR, i.e. *l*_2,1_-norm and trace norm) cause the sparse and similar connections cross each brain regions, which naturally incorporates the modularity prior of the estimated FBNs. The SLR FBN estimation model is given as follows:

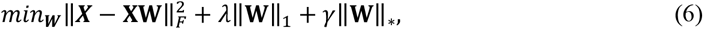

where ***X*** is the BOLD signal data, **W** is the estimated FBNs, *λ*‖**W**‖_1_ is the sparsity regularizer and *γ*‖**W**‖_*_ is the low-rank regularizer.

#### 2. 2. 3 Group Sparsity based FBN estimation

However, the above mentioned FBN estimation models are unable to deal with inter-subject variability problem since the FBN is estimated at an individual level, which will easily result in different network topological structures across subjects. To mitigate the effects of inter-subject variability, Wee et al proposed a group-constrained sparse linear regression model (Wee et al. 2014), which follow the idea of joint feature selection concept in group-lasso for regression problems (Yuan and Lin 2006). In particular, a group sparsity regularizer (GSR, i.e. *l*_2,1_-norm) is incorporated into the FBN estimation model. The GSR FBN estimation model is given as follows:

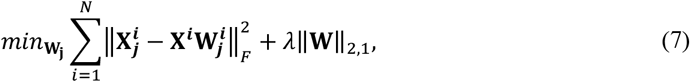

where 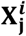 is the BOLD signal of *j*th ROI and *i*th participant, **X^*i*^** is the data matrix of *i* participant. 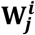 is the functional connections of the *j*th ROI and *i*th participant. *λ*‖**W**‖_2,1_ is the group sparsity regularizer. This minimize the inter-subject variability via an additional *l*_2_-norm penalization across all subjects than SR method. However, this methods may penalized too much for estimated FBNs, for example, a functional connection is collapse in MCIs and exist in NCs, if the number of NCs is larger than MCIs, the GSR method will tend to force this connect exists in MCIs, which will lose the discriminative information of the estimated FBNs.

#### 2. 2. 5 Our Methods

For easily incorporating the group constraint,

we first extent the existing matrix regularization framework to tensor form, which is defined as follows:

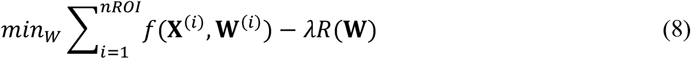

where **X** ∈ *R^nROI×T×n^* represent the input data, *nROI* is the number of predefined ROIs, *T* is the time length of the observed data, n is the number of participants. In particular, **X**^(*i*)^ represent the data matrix of the *i*th participant. **W** ∈ *R^nR0I×nR0I×n^* is the estimated FBNs, **W**^(*i*)^ represent the corresponding FBN of the ith participant. Obviously, both **X** and **W** are 3-dimensional tensor. Similar to the matrix regularization framework, in Eq. (8), 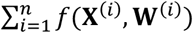is the data-fitting term and *R*(**W**) is the regularization term in tensor format.

The above mentioned *l*_2,1_-norm penalty excessive punished the estimated FBNs, which will lead to interference cross different groups in the data. For alleviating this issue, in this paper, based on the tensor regularization framework, we relax the *l*_2,1_-norm penalty and naturally introduce the tensor low-rank (TLR) regularizer for formulating the group similarity prior. The proposed tensor low-rank based FBN estimation can formulated as follows:

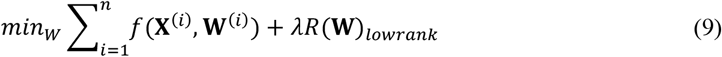

In Eq. (9), for the regularized terms, *R*(**W**)_*lowrank*_ indicates the rank of tensor **W**, which can be represented by number of non-zero elements in the eigenvalue of **W**. Unfortunately, the low-rank regularizer is non-convex with respect to **W** and it is NP-hard to solve. Thus, we relax it to trace-norm ‖**W**‖_*_, and obtain the following optimization model.

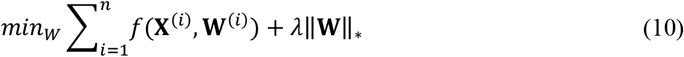

Here, due to its empirical effectiveness, we aim for capturing the second-order statistical structure of the observed fMRI data. In particular, we adopt SR as a testing platform, since the PC method suffers the cofounding effect. In particular, we use 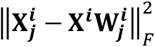 as the data-fitting term for formulating the inverse covariance structure (i.e., partial correlation) in the data, and adding a *l*_1_-norm penalty for encoding the sparse priors, and obtaining the following STLR optimization model.

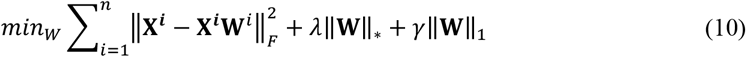

where *λ* and *γ* are regularized parameters used to control the balance among the three terms in the objective function. It should also be noted that the data fitting term can be designed as 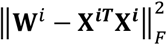 for capturing the full correlation statistics. In addition, when *γ* = 0, The proposed method reduces to the network learning model based on the traditional sparse regression FBN estimation method which gives in Eq. (4); when *λ* = 0, Eq. (8) reduces to the tensor low-rank representation FBN estimation method.

#### 2. 2. 4 Algorithm

For the reason of *l*_1_-norm and trace penalty exist, the proposed scheme is convex but non-differentiable, which leads to a nontrivial problem. Fortunately, several approaches are proposed for dealing with such issues (Donoho and Elad 2003, Meinshausen and Bühlmann 2006, Tomioka and Sugiyama 2009). In this paper, we select the proximal method (Combettes and Pesquet 2011) for solving the proposed optimal FBN estimation model, for the reason of its simplicity and efficiency. The details are given as follows:

Firstly, for the data-fitting term of STLR (i.e., 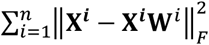), whose gradient w.r.t. *W^i^* are ∇_**w**_^i^*f*(**X^i^, W^i^**) = **X**^**i**^*T*^^**X^i^W^i^** − **X**^**i**^*T*^^**X^i^**. Therefore, for each iteration, we first update the **W**, according to the gradient descent criterion:

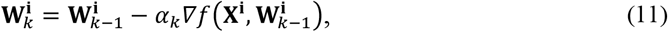

where *α_k_* denotes the step size of the gradient descent. The initial value of the step size *α_k_* is set to 0.001, and it will be adaptively updated in the following steps based on the line search scheme proposed by Nemirovski (NESTEROV 1983), according to the used SLEP toolbox3.

Secondly, for the regularization term ‖W‖_1_, according to the definition of proximal operator (Combettes and Pesquet 2011), the proximal operator of λ‖W‖_1_ is equivalent to the following soft thresholding operation on W,

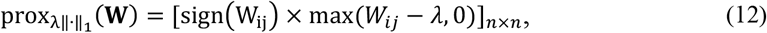

Similarly, the proximal operator of λ‖W‖_*_ corresponds to a shrinkage operation on the singular values of W, as follows.

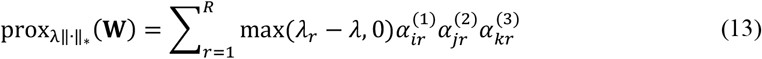

Here, *α_ir_, α_jr_, α_kr_* is a vector in a unit norm space, and the *λ_r_* is the corresponding eigenvalue based on the PARAFAC decomposition. Then, the final algorithm can be given as follows:

**Table 1.** The Algorithm for Estimating the FBN based on TLR

|  |
| --- |
| Input: $\mathbf{X}, \lambda$ |
| Output: $\mathbf{W}$ |
| Initialize $\mathbf{W}$ |
| while *not converged* |
| $\mathbf{W}_{k+1} = \text{gradient}(\mathbf{W});$ |
| $\mathbf{W}_{k+1} = \text{prox}_{\lambda\ \cdot\ _*}(\mathbf{W}_k);$ |
| $\mathbf{W}_{k+1} = \text{prox}_{\lambda\ \cdot\ _1}(\mathbf{W}_k);$ |
| end |

#### 2. 3. 3 Experimental Setting

After obtaining the FBNs of all subjects, the main task comes to use the constructed FBNs to train a classifier for identifying ASDs from NCs. Since the FBN matrix is symmetric, we just use its upper triangular elements as input features for classification. Even so, the dimensions of the features are still too high to train a classifier with good generalization, due to the limited training samples in this study. Therefore, we first conduct a feature filtering operation before training the classification. Specifically, the classification pipeline includes the following two main steps. In particular, we first estimated FBNs for each individual by PC^4^, SR, SLR, GSR and STLR, respectively. The estimated FBNs is shown in Fig. 1. After we obtain the estimated FBNs, the next task is how to use these connections for identifying MCIs from NCs. It should be noted that both the feature selection and classifier design have a big influence on the final accuracy (Wee et al. 2014). Therefore, in this paper, we only adopt the simplest feature selection method (t-test with p-value<0.01) and the most popular used SVM classifier with default parameter C=1 (Chang and Lin 2007), since our main focus is FBN estimation.

**Fig. 1.**
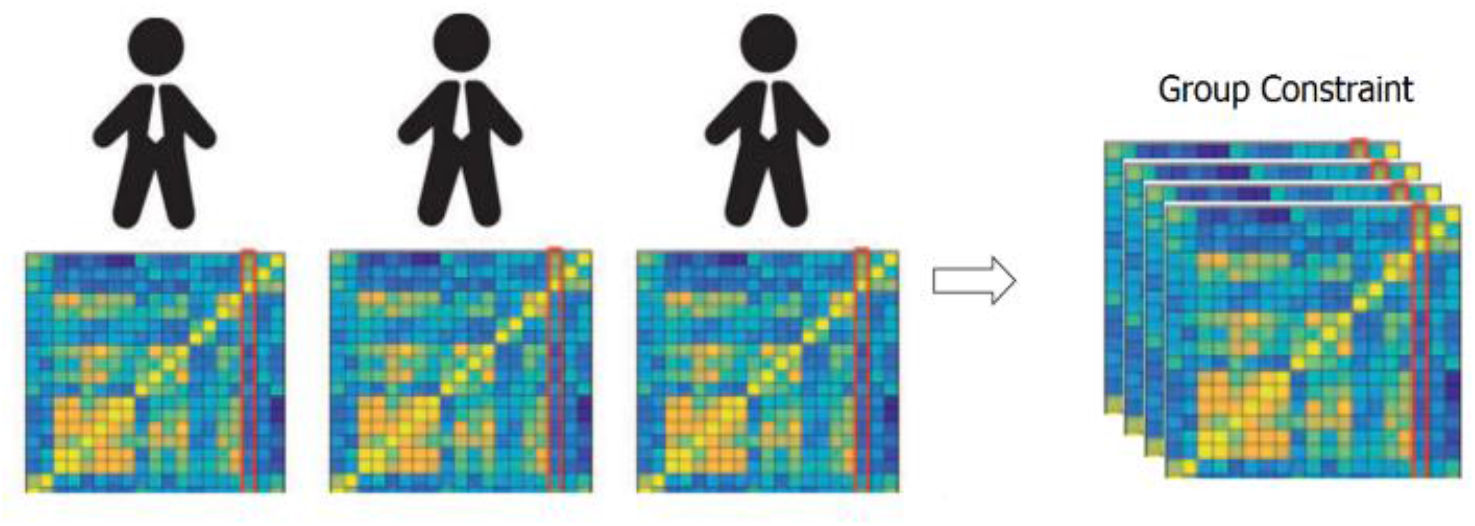
The motivation of the proposed tensor based FBN estimation model

Due to the small sample size, we use the leave one out (LOO) cross validation strategy to verify the performance of the methods, in which only one subject is left out for testing while the others are used to train the models and get the optimal parameters. For the choice of the optimal parameters, an inner LOO cross-validation is further conducted on the training data by grid-search strategy. More specifically, for the regularized parameter *λ*, the candidate values range in [2^−5^, 2^−4^,…, 2^4^, 2^5^]; for the hard threshold of PC_threshold_, we use 20 sparsity levels ranging in [5%, 10%,…, 95%, 100%]. For example, the 90% means that 10% of the weak edges are filtered out from the FBN.

## 3. Results

### 3.1 Network Visualization

For visual comparison of the FBN constructed by PC, SR, SLR, GSR and STLR methods, we first show the FBN adjacency matrices^5^ *W* constructed by different methods in **Fig. 1**.

It can be observed from Fig. 2 that the PC-based FBN (i.e. Fig.1 (a)) are quite different with the SR based FBNs (i.e. Fig.2 (b)-(f)), since it uses a different data-fitting term (i.e., the first term in Equation (5)). Moreover, the topology of the FBN estimated by SLR is similar to that of STLR, because (1) both methods employ the same data-fitting term, and (2) the low-rank and sparse regularity behind SLR (i.e., the trace norm in matrix scheme) is based on the result of STLR (i.e., the trace norm in tensor scheme).

**Fig. 2.**
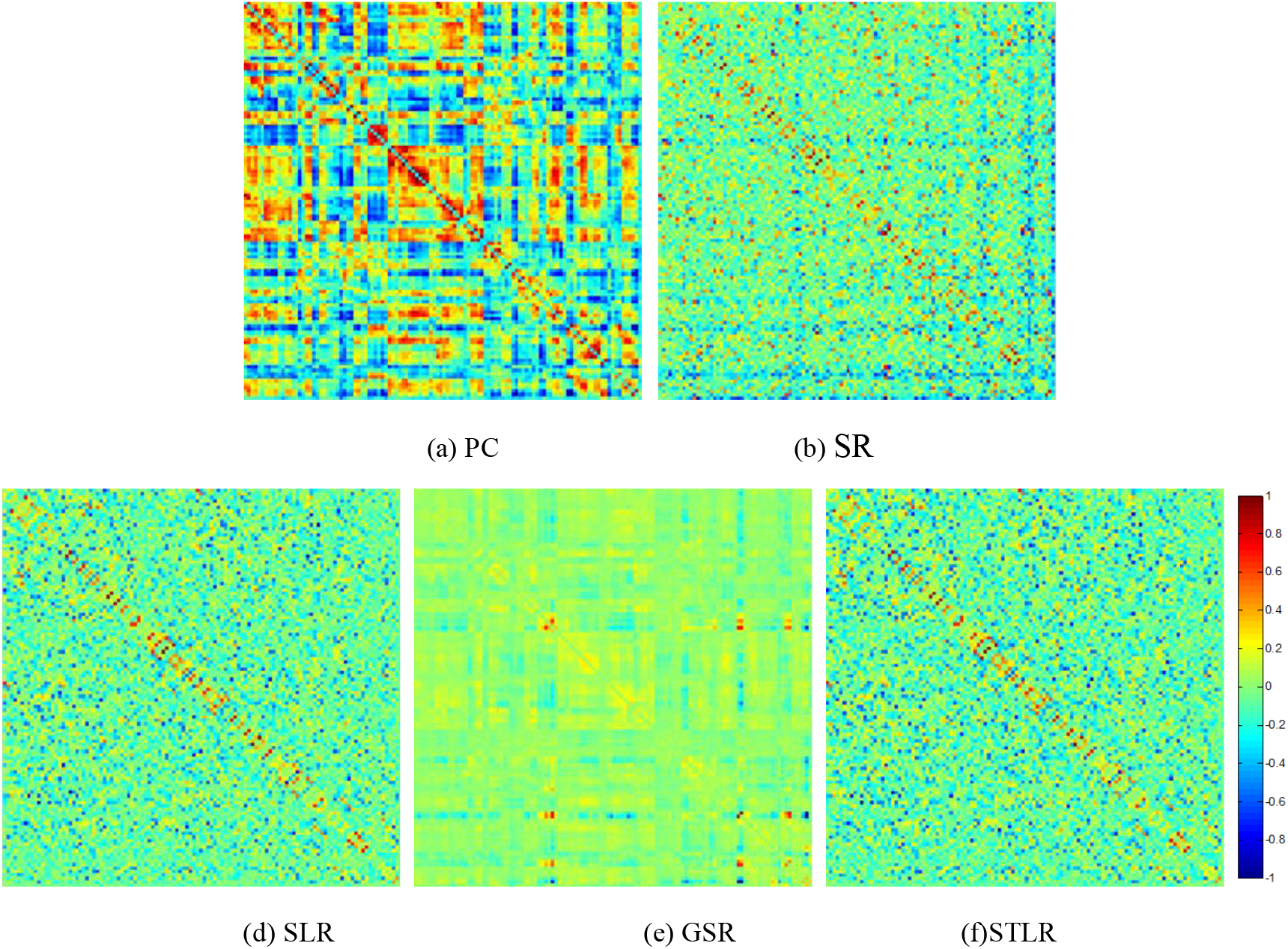
The FBN adjacency matrices of a certain subject, constructed by different methods.

### 3.2. MCI Identification

The MCI vs NC classification results on ADNI dataset are given in **Table 2**. The proposed STLR method achieves the best accuracy in this experiment. In addition, the results of SLR and GSR are also provided in **Table 2** as a reference.

**Table 4.** Classification performance corresponding to different FBN estimation methods on ADNI dataset.

| Method | Accuracy | Sensitivity | Specificity |
| --- | --- | --- | --- |
| PC | 67.15 | 72.06 | 62.32 |
| SR | 78.10 | 79.41 | 76.81 |
| SLR | 80.29 | 80.88 | 79.71 |
| GSR | 83.21 | 88.24 | 78.26 |
| STLR | **91.97** | **92.65** | **91.30** |

A set of quantitative measurements, including accuracy, sensitivity and specificity, are used to evaluate the classification performance of four different methods (PC, SR, SLR, GSR and STLR). The mathematical definition of these three measures are given as follows:

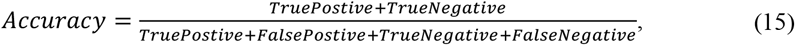

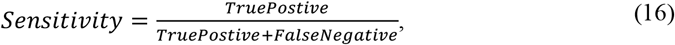

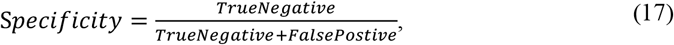

Here, *TruePositive* is the number of the positive subjects that are correctly classified in the ASD indentification task. Similarly, *TrueNegative*, *FalsePostive* and *FalseNegative* are the numbers of their corresponding subjects, respectively.

### 3.3. Most Discriminant Connections and Brain Regions

For discovering the biological meaningful biomarkers, we also provided the most significant connections estimated by STLR between MCIs and NCs. We firstly identified the most significant connections (p-value<0.01) and projected them into the corresponding subnetworks (Fig.3). We found that the most discriminative network connections were distributed mainly in the default mode network (DMN), frontoparietal task control network, salience network and visual network.

**Fig. 3.**
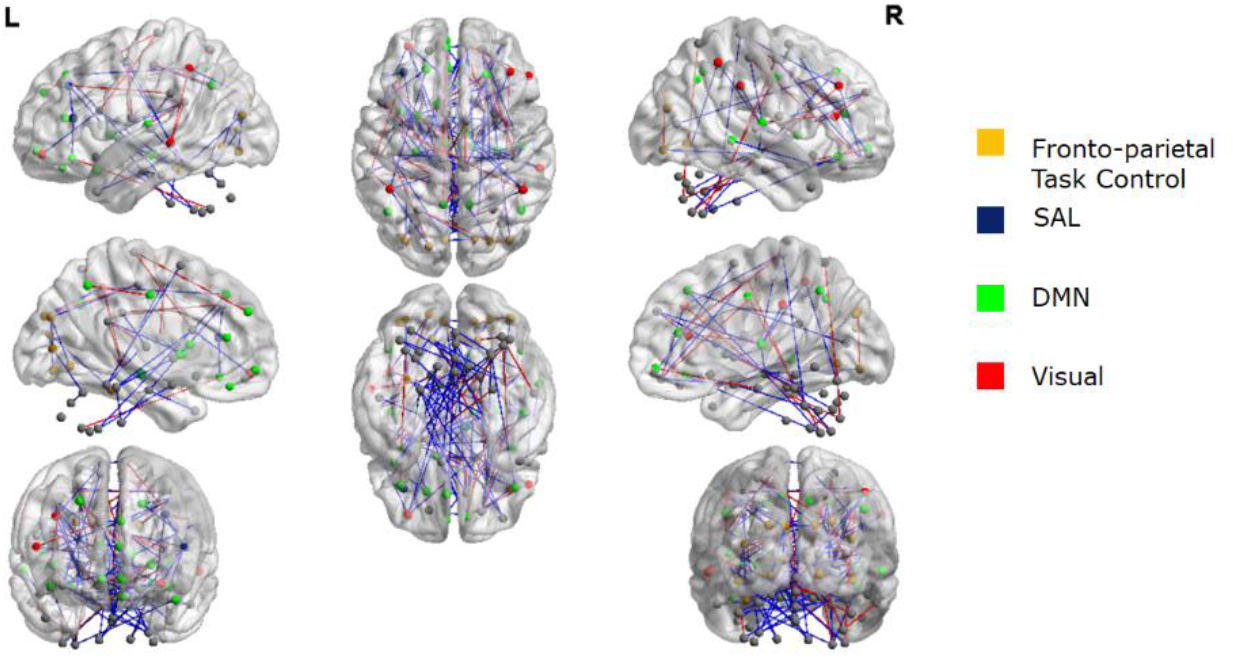
The most significant functional connections mapped on the ICBM 152 template using the BrainNetViewer package (http://nitrc.org/projects/bnv/). The details are: yellow, frontoparietal task control network; blue, salience network; green, default mode network; red, visual network.

Meanwhile, we projected these significant functional connections into the corresponding brain regions. The most 130 significant edges with p-value<0.01 is depicted in Fig. 4 with the width of each arc represents the weight of the connection between two end points (i.e., brain regions). These brain regions were considered as the extremely predominant areas with largest number of the discriminative connections based on the proposed FBN construction model for discriminating MCI participants from NCs. The location of these ROIs was labeled according to AAL atlas.

**Fig. 4.**
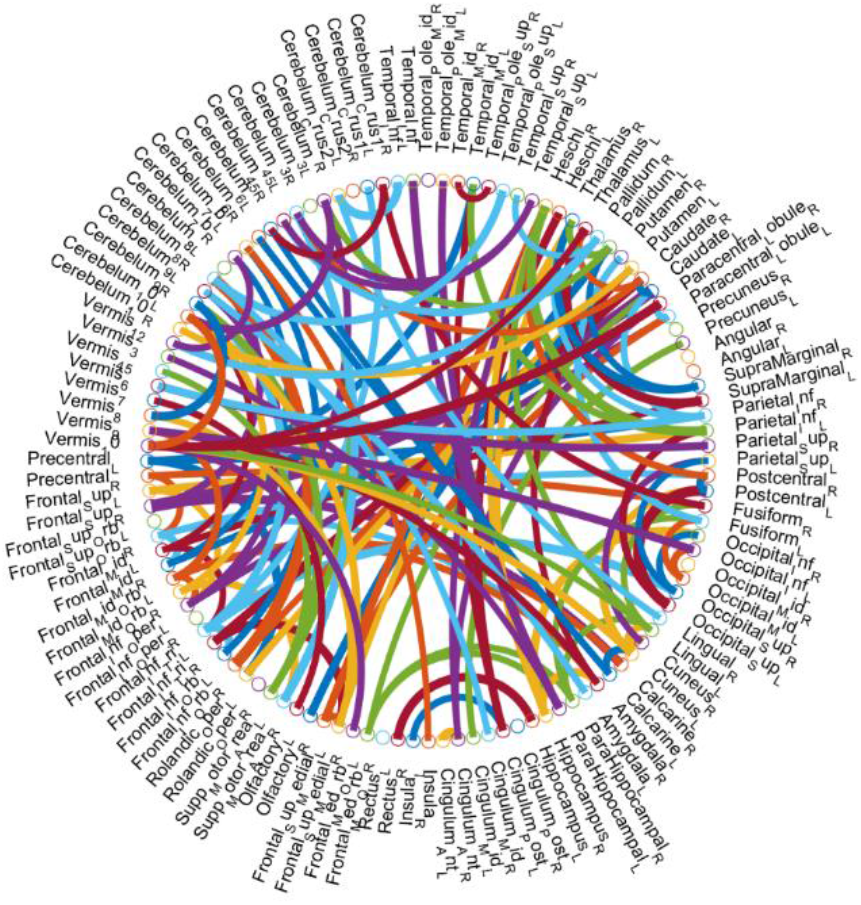
The most discriminative connections, selected by t-test (p<0.01), between MCI and NC for 116 ROIs of AAL template,. This figure is created by a Matlab function, circularGraph, shared by Paul Kassebaum. http://www.mathworks.com/matlabcentral/fileexchange/48576-circulargraph

**Fig. 4.**
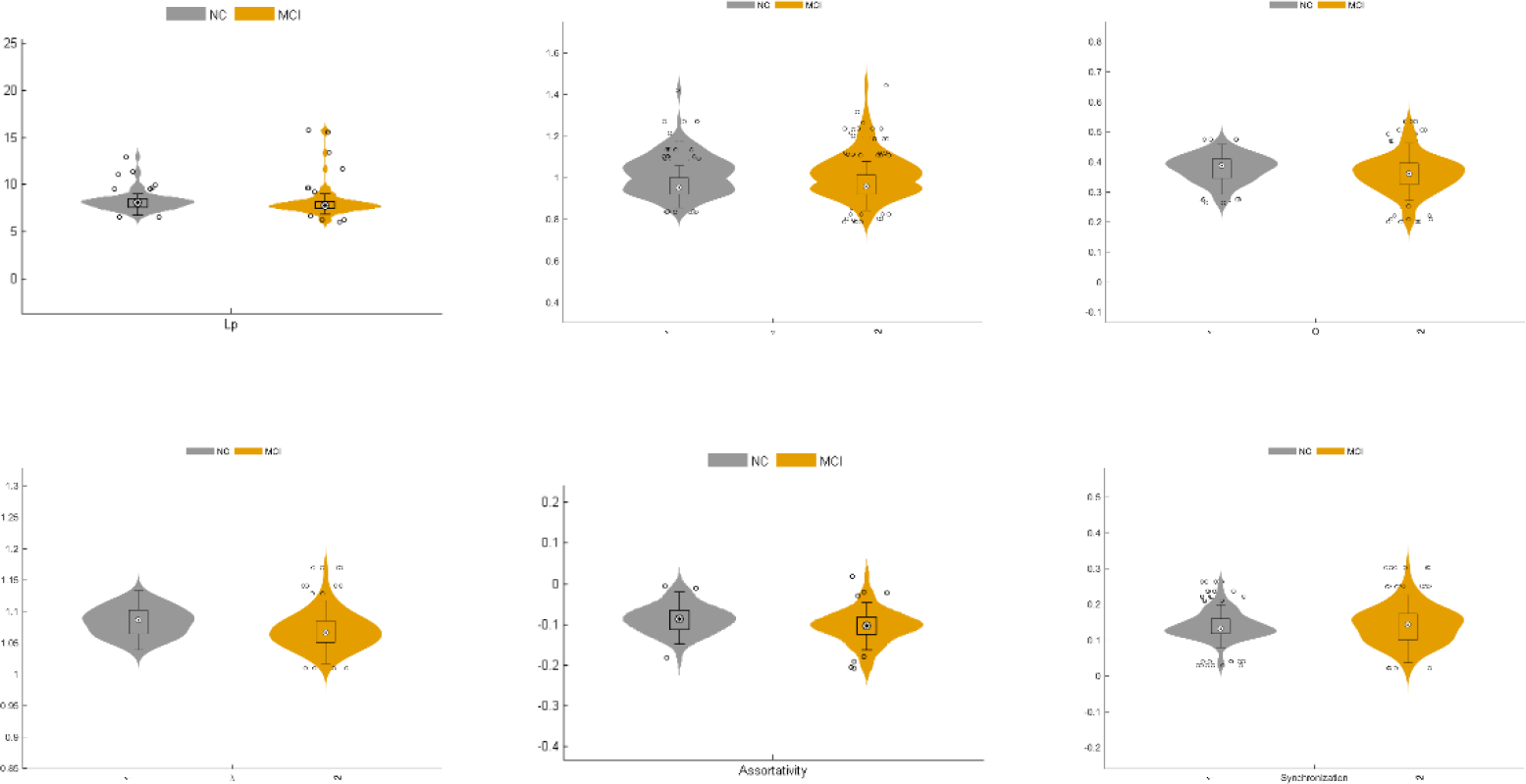
The Global graph metrics of the FBNs in MCI and NC groups.

As shown in Table 5, the most discriminative brain regions between MCI and NCs were distributed mainly in the thalamus, middle temporal gyrus, hippocampus, parahippocampal gyrus, inferior parietal which corresponded to the subcortical network, DMN, dorsal attention network, and fronto-parietal task control.

**Table 5.** Top-20 brain regions (without cerebellum region) with largest number of the discriminative connections

| AAL Number | Corresponding brain regions | Sub-networks |
| --- | --- | --- |
| 77 | Thalamus_L | Subcortical network |
| 85 | Temporal_Mid_L | Dorsal attention network |
| 6 | Frontal_Sup_Orb_R | Default mode network |
| 9 | Frontal_Mid_Orb_L | Default mode network |
| 38 | Hippocampus_R | Default mode network |
| 39 | ParaHippocampal_L | Default mode network |
| 61 | Parietal_Inf_L | Dorsal attention network |
| 62 | Parietal_Inf_R | Dorsal attention network |
| 70 | Paracentral_Lobule_R | Sensory/somatomotor hand |
| 71 | Caudate_L | Fronto-parietal task control |
| 72 | Caudate_R | Fronto-parietal task control |
| 75 | Pallidum_L | Subcortical network |
| 11 | Frontal_Inf_Oper_L | Executive control network |
| 13 | Frontal_Inf_Tri_L | Executive control network |
| 24 | Frontal_Sup_Medial_R | Fronto-parietal task control |
| 42 | Amygdala_R | Subcortical network |
| 45 | Cuneus_L | Visual network |
| 47 | Lingual_L | Default mode network |
| 73 | Putamen_L | Salience network |
| 78 | Thalamus_R | Subcortical network |

AAL: the automated anatomical labeling atlas.

### 3.4. Altered topological properties of functional networks in MCI patients

Based on the estimated FBNs by STLR, several global graph theory metrics, including clustering coefficients (C_p_), shortest path length (L_p_), normalized clustering coefficient (γ), normalized characteristic path length (λ) small-world (σ) and global efficiency (E_global_), were calculated to uncover the topological properties of functional networks in MCI and NC groups. As expected, both two groups fit γ =C_p_^real^ / C_p_^rand^ > 1, λ=L_p_^real^ / L_p_^rand^ ≈ 1 and σ= γ /λ> 1. Thus, the functional networks of MCI patients and NCs showed the topological attributes of small-world (Watts and Strogatz 1998). This means the brain networks of the two groups maintain a complex and efficient neural architecture that optimizes the balance between local specialization and global integration (Achard 2007, Sporns and Zwi 2004, Sporns 2012). Further comparison suggested that the small-world σ value of MCI patients was lower than that of NCs, which indicated the disruption of “economic small-world” (i.e., reduction of the segregation and integration functions of effective information in the brain network). Furthermore, we found the C_p_ value and modularity (Q value) in MCI patients were lower than that in NC groups significantly (P<0.01). These changes of Cp and modularity suggest the reduction of network segregation in local information processing in MCI patients. Although there was no significant difference between MCI and NCs in L_p_ and E_global_, the lower values of these two global topological attributes in MCI indicated the decreased network integration.

According to the definition of “hubs”, we identified hub nodes in MCI patients and NCs. As shown in table 6, the common hub regions of MCI and NCs were located mainly in the bilateral superior temporal gyrus, bilateral heschl gyrus, right middle frontal gyrus and left angular gyrus. Most of them mainly distributed in the DMN, Auditory network, fronto-parietal task control network and dorsal attention network. Moreover, it is notable that some hub nodes were present only in MCI patients and absent in HCs: right middle temporal gyrus, left middle frontal gyrus. Simultaneously, there were also some hub nodes in HCs but not in MCI patients. They were located on the right inferior parietal and right middle frontal gyrus. These discriminative brain regions between MCI and NCs were distributed mainly in DMN, fronto-parietal task control and dorsal attention network. The differences of subnetworks and corresponding brain regions play an important role in the differential diagnosis of MCI and NC.

**Table 6.** Hubs in MCI and NCs defined with the degree

|  | AAL Number | Corresponding brain regions | Sub-networks |
| --- | --- | --- | --- |
| MCI | 88 | Temporal_Pole_Mid_R | Default mode network |
|  | 79 | Heschl_L | Auditory network |
|  | 10 | Frontal_Mid_Orb_R | Default mode network |
|  | 80 | Heschl_R | Auditory network |
|  | 81 | Temporal_Sup_L | Auditory network |
|  | 65 | Angular_L | Default mode network |
|  | 87 | Temporal_Pole_Mid_L | Default mode network |
|  | 9 | Frontal_Mid_Orb_L | Fronto-parietal task control |
|  | 83 | Temporal_Pole_Sup_L | Auditory network |
|  | 84 | Temporal_Pole_Sup_R | Auditory network |
| NC | 79 | Heschl_L | Auditory network |
|  | 81 | Temporal_Sup_L | Auditory network |
|  | 65 | Angular_L | Default mode network |
|  | 66 | Angular_R | Default mode network |
|  | 87 | Temporal_Pole_Mid_L | Default mode network |
|  | 10 | Frontal_Mid_Orb_R | Default mode network |
|  | 80 | Heschl_R | Auditory network |
|  | 62 | Parietal_Inf_R | Dorsal attention network |
|  | 84 | Temporal_Pole_Sup_R | Auditory network |
|  | 83 | Temporal_Pole_Sup_L | Auditory network |

AAL: the automated anatomical labeling atlas.

## 4. Discussion

The human brain is the most complex systems in the world. In order to ensure an efficient interaction of information in the brain, the FBN should has more “structures” than just sparsity (Smith et al. 2011b, Sporns 2011). In this work, we incorporated a tensor low rank regularizer for modeling the group similarity priors of the estimated FBNS. The proposed models were verified on the ADNI dataset for MCI *vs* NC classification. Based on the results, we give the following brief discussion.

1. The accuracy of the STLR-based methods outperforms the baseline and the states-of-art method on our used dataset. A possible reason is that the STLR scheme naturally incorporate more information from inter-group subjects, and thus can gain more clearly or discriminative FBNs. It should also be noted that the proposed scheme is a flexible module, meaning that, besides the SR-based models, it can be easily adopted on other FBN estimation models such as PC-based network, Bayesian network or Granger Causal based network. Also, we can incorporate some other useful priors such as modularity, scale-free into the Tensor based FBN estimation models.
2. The most discriminative functional connections and the corresponding predominating brain regions. By projecting brain regions with significant differences of functional connectivity and graph theory metrics in the brain network to subnetworks, we found that the differences between MCI patients and NCs were distributed mainly in the DMN, dorsal attention network, frontoparietal task network, executive control network and auditory network. Especially, the DMN had the most significant discriminative ability. The changes of these subnetworks were consistent with the results of previous studies on cognitive function (i.e., spatial attention (Rolle et al. 2017), executive function (Liao et al. 2019)and auditory function (Bi et al. 2018)) corresponding to subnetworks in MCI patients. Moreover, the DMN has been regarded to be the core part of functional center (Liu et al. 2019), which is involved in episodic memory and has been considered as the major cognitive domain impaired in the early stage of AD (Eyler et al. 2019). Furthermore, in our study, except for validating the discriminative ability of the DMN for MCI identification, we located the predominant brain regions (i.e., the thalamus, middle temporal gyrus, hippocampus, parahippocampal gyrus, inferior parietal and middle frontal gyrus). These results could be beneficial for the early, accurate diagnosis of MCI.
3. Altered pattern of the brain network connectome in MCI. In our study, firstly, we found that MCI patients and NCs fitted the small-world attribute in the global topological property. That is, the brain network of MCI and NC groups conform to “economic small-world” which can rapid, real-time information processing across separate brain regions to maximize efficiency with minimal cost and to render resilience against pathological attacks (Liao, Vasilakos and He 2017, Sporns and Zwi 2004, Sporns 2012). Statistical analysis suggested that the value of small-world σ in MCI patients was lower than that in NCs, which indicated disruption of brain network integration and segregation. This result of small-world in MCI is consistent with some previous research (Yu et al. 2018). Moreover, the significantly decreased value of C_p_ and modularity in MCI further verified the reduction of functional segregation of brain network. A lower value of C_p_ and Q value of modularity suggest the less concentrated clustering of local connections and weaker capacity for specialized processing of within densely interconnected groups of brain regions in MCI (Rubinov and Sporns 2010).

However, since the proposed scheme is a simple try for modelling the group similarity prior, there are several limitations in the proposed methods that need to be improved in the future work.

1. In this paper, we only provide a simply verification for validating the effectiveness of the TL scheme, without considering other factors (e.g. the selection of Atlas and data preprocessing). Therefore, we simply adopt the commonly used AAL atlas to define ROI. In the future, we would like to consider the functional template (e.g. Power264) for alleviating this issue.
2. In this paper, we only use the tensor low-rank module to formulate the group similarity prior. In fact, the brain has a very complex structure, and the group similarity can also be formulated into another format. Therefore, in the following, we will use more abundant prior information or topology structure to construct appropriate regular terms and further improve the current group-constraint model.
3. The global graph theory metrics (i.e., C_p_, L_p_, small-world) were mainly discussed in our study, while nodal and other graph theory metrics could also be used to describe the complex topological mechanism of brain networks. In the following research, more graph theory metrics, such as nodal shortest path length, local efficiency and participant coefficient of modularity can be used to elaborate more specific topological properties of local brain network.

## 5. Conclusion

In fact, the pattern of human brain still needed a deep exploration, therefore, how to better describe the brain is still a challenging and meaningful. In sprite by the fact that the group similarity constraint of FBN cross group, we naturally introduce the tensor low-rank regularer based FBN estimation scheme. In particular, we use the PARAFAC decomposition for capturing the FBNs with low rank topologies. In the end, we put the estimated FBNs into the classification task. The result illustrate that the introduction of the group similarity constraint can effectively improve the performance of the baseline method. The post hoc analysis of the graph theory metrics further shown more biologically meaningful functional brain connections obtained by our proposed method.

## AUTHOR CONTRIBUTIONS

All authors developed Proposed algorithm, architecture. Wei-kai Li and Rui Li designed the evaluation experiments. Xin Gao preprocessed the fMRI. Xiaowen Xu analyzed and interpreted the results of data. All authors contributed to preparation of the article, figures, and charts.

## ACKNOWLEDGMENTS

This work was partly supported by Scientific and Technological Research Program of Chongqing Municipal Education Commission (KJ1500501, KJ1600512, Kj1600518), and Shanghai Municipal Planning Commission of Science and Research Fund (201740010).

## Footnotes

1 http://adni.loni.ucla.edu

2 http://www.fil.ion.ucl.ac.uk.spm

3 http://www.yelab.net/software/SLEP

4 In order to improve the flexibility of PC and conduct fair comparison, we introduce a hard-thresholding parameter in PC by reducing a proportion of weak connections.

5 The adjacency matrix is an algebraic expression of a graph (or network). The elements of the matrix indicate the connection strength of the node pairs in the graph. Here, for the convenience of comparison among different methods, all the weights are normalized to the interval [−1 1]

